# Updated Guidance for Communicating PFAS Identification Confidence with Ion Mobility Spectrometry

**DOI:** 10.1101/2025.01.27.634925

**Authors:** Anna K. Boatman, Jessie R. Chappel, Kaylie I. Kirkwood-Donelson, Jonathon F. Fleming, David M. Reif, Emma L. Schymanski, Julia E. Rager, Erin S. Baker

**Affiliations:** Department of Chemistry, University of North Carolina at Chapel Hill, Chapel Hill, North Carolina 27514, USA; Department of Environmental Sciences and Engineering, Gillings School of Global Public Health, The University of North Carolina at Chapel Hill, Chapel Hill, North Carolina 27514, USA; The Institute for Environmental Health Solutions, Gillings School of Global Public Health, The University of North Carolina at Chapel Hill, Chapel Hill, North Carolina 27514, USA; Curriculum in Toxicology and Environmental Medicine, School of Medicine, University of North Carolina, Chapel Hill, North Carolina 27514, USA; Immunity, Inflammation, and Disease Laboratory, National Institute of Environmental Health Sciences, Durham, North Carolina 27709, USA; Predictive Toxicology Branch, Division of Translational Toxicology, National Institute of Environmental Health Sciences, Durham, North Carolina 27713, USA; Luxembourg Centre for Systems Biomedicine (LCSB), University of Luxembourg, 6 Avenue du Swing, L-4367, Belvaux, Luxembourg

**Keywords:** ion mobility spectrometry (IMS), non-targeted analysis (NTA), high-resolution mass spectrometry (HRMS), identification confidence, per- and polyfluoroalkyl substances (PFAS)

## Abstract

Over the last decade, global contamination from per- and polyfluoroalkyl substances (PFAS) has become apparent due to their detection in countless matrices worldwide, from consumer products to human blood to drinking water. As researchers implement non-targeted analyses (NTA) to more fully understand the PFAS present in the environment and human bodies, clear guidance is needed for consistent and objective reporting of the identified molecules. Confidence levels for small molecules analyzed and identified with high-resolution mass spectrometry (HRMS) have existed since 2014; however, unification of currently-used levels and improved guidance for their application is needed due to inconsistencies in reporting and continuing innovations in analytical methods. Here, we (i) investigate current practices for confidence level reporting of PFAS identified with liquid chromatography (LC), gas chromatography (GC), and/or ion mobility spectrometry (IMS) coupled with HRMS and (ii) propose a simple, unified confidence level guidance that incorporates both PFAS-specific attributes and IMS collision cross section (CCS) values.

**Synopsis:** Unified and simplified requirements to guide confidence level assignment in non-targeted PFAS identification efforts with ion mobility spectrometry.

**Graphical Abstract:** For Table of Contents only

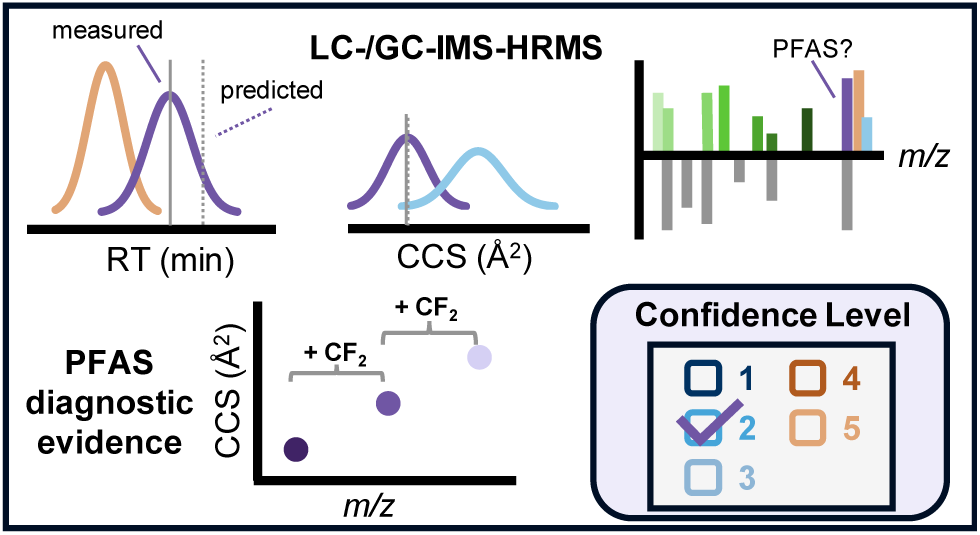

## Introduction

Per- and polyfluoroalkyl substances (PFAS) are a large and growing class of anthropogenic fluorinated chemicals^1–3^, with estimates ranging from 4,700 chemicals on the widely cited National Institute of Standards and Technology (NIST) PFAS Suspect List^4^ to over 7 million PFAS in PubChem.^5^ As PFAS production, use, and monitoring expands, these numbers are likely to increase. To date, researchers have typically focused on a limited number of PFAS due to the low availability of analytical standards, but the increased use of non-targeted analyses (NTA) with high resolution mass spectrometry (HRMS) has enabled broader monitoring and novel chemical discovery.^1–3^ Similarly, the use of analytical techniques beyond gas or liquid chromatography (GC or LC) coupled to HRMS is expanding. For example, numerous studies cite ion mobility spectrometry (IMS) as a promising technology for PFAS research due to its ability to separate PFAS from biomolecules,^1, 2, 6, 7^ and its adoption is already growing.

When reporting NTA results, it is critical to assign a confidence level to each identified molecule.^8^ Confidence levels provide context for chemical identifications and are important for transparent reporting of results,^9^ interpretation of exposure and toxicity data,^10^ and appropriate response in the event of an environmental release of unknown chemicals.^11^ Schymanski *et al.*^8^ first defined confidence levels for small molecule identifications made using HRMS in 2014 and others have subsequently adapted these levels. For example, Celma *et al*.^12^ incorporated CCS values into the 2014 levels, and Charbonnet *et al.*^3^ defined a PFAS-specific system, adding additional sublevels to the 2014 levels using PFAS-specific attributes. However, manual level assignments following any of these schemes are often subjective and prone to unintended user bias (often in the form of overestimation), because lower confidence identifications are often perceived as “worse” than higher confidence ones. Additionally, little guidance is available for what to do when some parameters do not meet the defined requirements.^12–14^ Even our own extensive experience with PFAS NTA using LC-IMS-HRMS has frequently generated situations where confidence levels are not defined by any existing guidance.^15, 16^ Further, there are vast and growing numbers of PFAS candidates, and a rapidly increasing number of researchers performing PFAS NTA on different analytical platforms including GC-HRMS, LC-HRMS, LC-IMS-HRMS^17–21^, strengthening the need for simple and objective confidence level assignment to support PFAS NTA. Here, we investigate the current reporting of PFAS NTA identifications, address identified gaps by clarifying and unifying existing guidance, and introduce a simplified checklist to help standardize confidence level assignments for PFAS identified using LC-IMS-HRMS.

## Materials and Methods

To investigate current practices for confidence level reporting of PFAS identified in NTA using HRMS, information was aggregated across our team’s extensive experience in this field as well as a dedicated literature review examining recently published papers worldwide. Our team’s experience includes the analysis of hundreds of analytical PFAS standards and numerous environmental matrices (water, serum, plasma, whole blood, tissue, etc.).^7, 15, 16, 22–24^ For the literature review, a search was performed in PubMed (RRID: SCR_004846) on 30 September 2024 using the following search query: ((PFAS[Title/Abstract]) OR (polyfluor*[Title/Abstract]) OR (perfluor*[Title/Abstract])) AND ((non-target*[Title/Abstract]) OR (nontarget*[Title/Abstract]) OR (untarget*[Title/Abstract])). A total of 426 publications from as early as 1997 were identified through this query, with the 94 studies published in 2023 selected for manual review to assess research trends in more detail. Inclusion in the in-depth analysis required articles specifically use HRMS for PFAS analysis, yielding 38 studies in total from 2023.^25–62^ Information from these articles was then summarized, including platform type, chemical identification reporting approach, matrix analyzed, number of PFAS identified, and confidence level scheme cited (see **Table S1**). This information was then used to help create the guidance criteria for assigning confidence levels to PFAS identified using LC-IMS-HRMS by understanding gaps in literature and practices which need to be considered in the guidance.

## Results and Discussion

### Assessing Current Practices for Confidence Levels in PFAS NTA

Our literature search identified 38 papers published in 2023 using HRMS for PFAS NTA worldwide (**Table S1**). The most common platforms were LC-Orbitrap and LC-QTOF (see **Table S2** for full breakdown of instruments and experiment types). One study used Fourier transform ion cyclotron resonance (FTICR) and no chromatography, and one study did not specify the type of HRMS instrument used. This illustrates the wide range of platforms available for PFAS NTA and the lack of transparency in reporting. Some studies analyzed PFAS exclusively, while others evaluated a broader range of pollutants that also included PFAS. The types of experiments conducted were diverse, including environmental monitoring (searching for unknown molecules in samples collected from subjects or nature), remediation studies (evaluating PFAS breakdown via intentional degradation technology treatment), industrial analysis (investigating composition of consumer products), and more. A variety of matrices were analyzed, including water (most common), blood, soil, air and analytical standards. Studies identified a wide range of PFAS (0 to 102 PFAS per study; **Figure S1**), and interestingly, more than a quarter of these studies (12) did not report confidence levels (**Figure 1**). Among the 26 studies that reported confidence levels, the 2014 levels were the most cited for PFAS identifications. Additionally, we observed a wide distribution of confidence level assignments, with Level 2 being the most frequently assigned (**Figure 2**). Previous guidance by Schymanski *et al*.^8^ and Celma *et al*.^12^ stated that Level 2b (probable structure by diagnostic evidence) identifications are usually rare;^8, 12^ however, we found that was not the case with PFAS NTA. In fact, 8 of the 10 studies from the literature search that used sublevels for Level 2 reported Level 2b and/or 2c identifications (range: 5 to 87 PFAS at Level 2b/c PFAS.^25, 32, 35, 37, 38, 54, 55, 61^ Guidance from Charbonnet *et al.*^3^ defines Level 2b [probable by diagnostic fragmentation evidence] and 2c [probable by diagnostic homologue evidence] as equivalent confidence). An in-depth evaluation of the methods showed that the confidence level guidelines were applied inconsistently (three such examples are shown in **Figure S2**). Examples include reported mass errors exceeding author-defined thresholds or assigned levels that contradict established guidance. For example, in some cases Level 2b was assigned to structures with explicitly noted potential isomers. However, one exact structure is required for Level 2b, and the possibility of isomers points to Level 3 (tentative candidates per Schymanski *et al.* or Celma *et al.* guidance^8, 12^, or positional isomer candidates per Charbonnet *et al.* guidance^3^). These inconsistencies complicate cross-study comparisons even across identical analytical platforms and call into question the accuracy of reported confidence levels. These inconsistencies not only hinder the interpretation of published data but also impede comparison of IMS to non-IMS results, and underscore the need for more objective, standardized confidence level assignments. Clearly, there is an opportunity to unify existing confidence level frameworks in a way that enables clear, simple and more comparable reporting as IMS is increasingly adopted in PFAS NTA.

**Figure 1.**
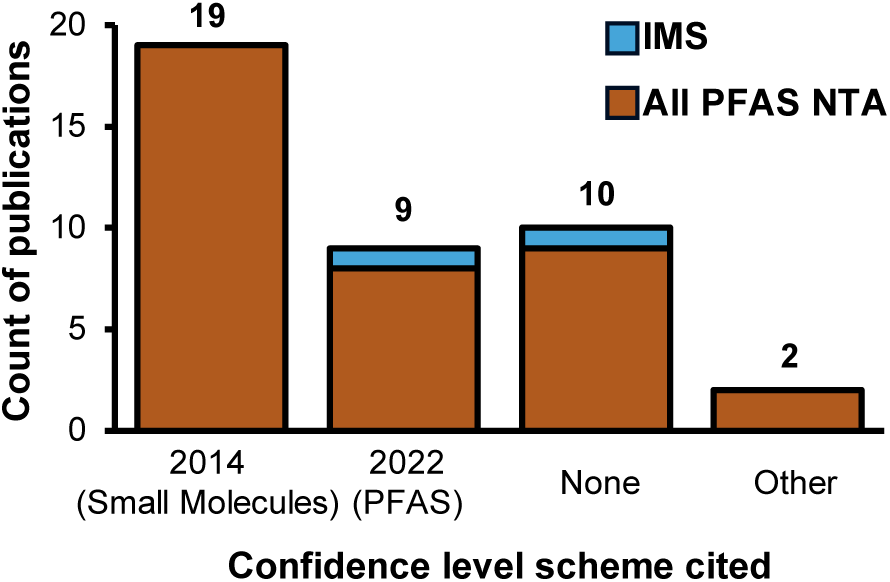
Confidence level scheme cited in the 38 papers from 2023. 2014 (Small Molecules): cited Schymanski *et al*. scale for small molecules. 2022 (PFAS): cited Charbonnet *et al*. scale for PFAS. None: no confidence levels were reported. Other: cited different confidence levels that do not use a 1-5 scale.

**Figure 2.**
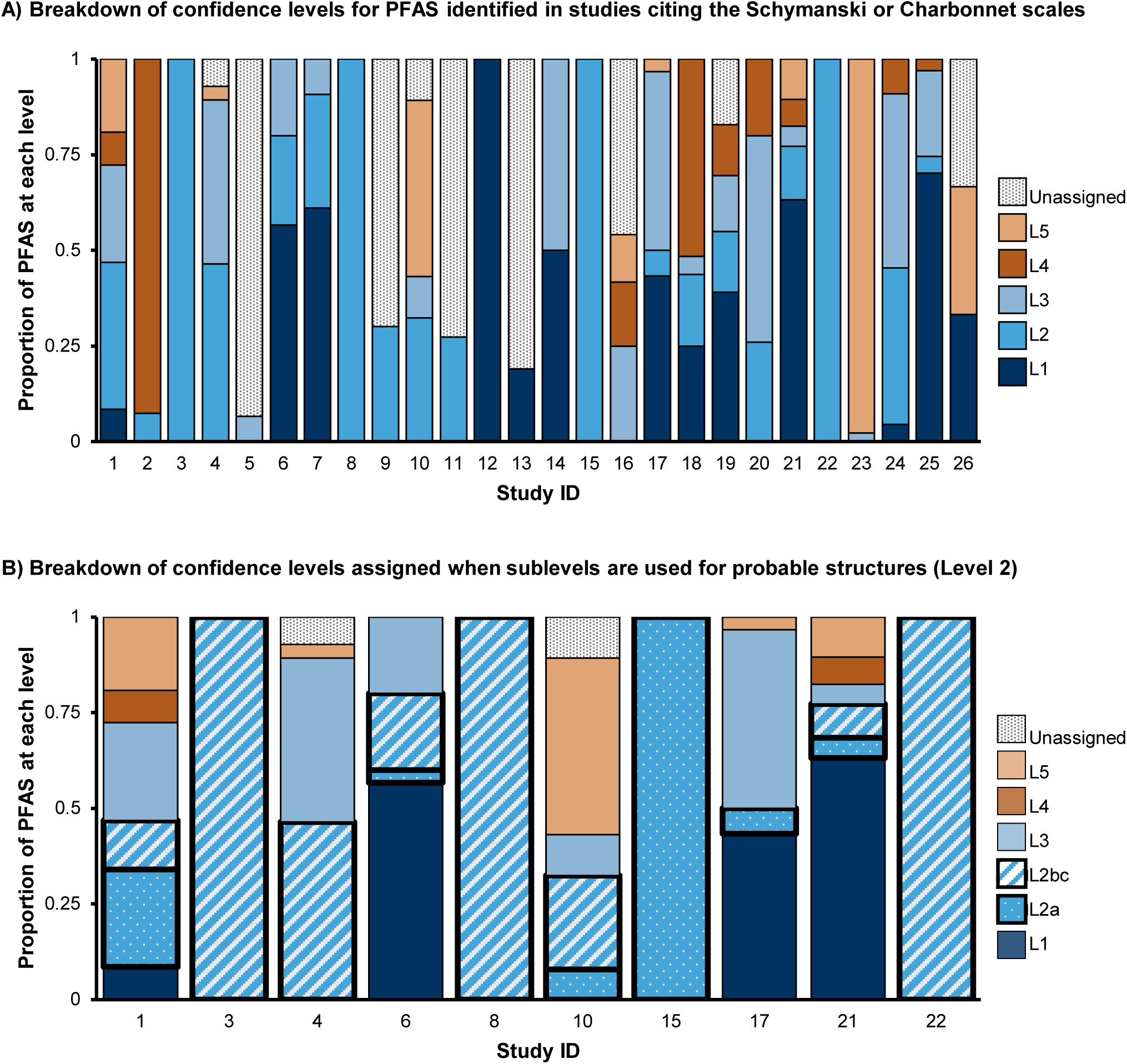
Distribution of confidence levels among the 26 papers that identified > 0 PFAS and cited Schymanski or Charbonnet scales. **A)** Percent of PFAS identified at each level by study. In several instances not all PFAS identified were assigned a confidence level, e.g. Study 9 reported 93 PFAS identified, assigned Level 2 to 28 of them, and did not assign confidence levels to the remainder. **B)** Breakdown of the 10 papers that assigned sublevels to Level 2 probable structures. 9 papers did not assign sublevels, and 8 papers did not report any Level 2 detections. Level 2 identifications are shown with thicker outlines and pattern fill (dots for 2a, diagonal stripes for 2b and 2c). The Charbonnet scale defines 2b and 2c as equivalent confidence, so these levels are combined to facilitate comparison to the Schymanski scale.

To understand potential improvement areas for confidence levels, the 38 NTA studies were evaluated further. A lack of universal standards for acceptable NTA performance^3, 13, 14, 63^ has been noted elsewhere and was evident in the literature evaluation. For instance, mass tolerances were sometimes not defined or set as high as 20 ppm for HRMS data. Automation is one potential solution for these inconsistencies, where confidence levels would be assigned objectively based on predefined criteria for mass accuracy and other dimensions. For automation to function, discrete criteria must be provided. Existing scales have attempted to define criteria for mass, CCS, and RT accuracy. However, given the wide variety of analytical techniques and modeling tools already used and being developed for PFAS NTA, universally suitable acceptance tolerances cannot yet be defined. For example, in other literature beyond the 38 studies evaluated here, experimental CCS values can be highly reproducible (with reported reproducibility of ≤ 0.30% in some cases),^64^ while predicted CCS values are more variable, with Oone study noting machine learning tools with errors up to 8%.^23^ All methods have utility depending on the experimental context, and defining an explicit tolerance to apply universally would be detrimental. Thus, we faced a challenge in unifying objective tolerance criteria. We propose a potential way forward of using an instrument- and method-based set of recommended tolerances, as discussed further below.

### Addressing Gaps in Existing Confidence Level Assignments

To address the current challenges in communicating confidence for PFAS identified using LC-IMS-HRMS our first step was to unify and simplify the existing confidence level guidance for small molecules^8^, IMS^12^, and PFAS^3^. An overarching observation of our literature search was an inconsistent application of current guidance, leading us to conclude that simplification and clarification of existing guidance was necessary. The updated requirements are summarized in a checklist^65^ (**Figure 3**), described in detail in the following section, and provided as an extended printable document in the **Supplemental Checklist**. One key change is the removal of sublevels. Existing scales distinguish between a probable structure based on a library match (Level 2a) and one based on diagnostic evidence if no library entry exists (Level 2b). Charbonnet *et al.* also defined a Level 2c, defined as equivalent to Level 2b for probable structures elucidated using PFAS homologous series. However, in our experience analyzing PFAS using LC-IMS-HRMS, we have detected PFAS that exist in spectral libraries but not CCS libraries. We have also detected PFAS that match our in-house CCS library, fall exactly on CCS vs *m/z* and RT vs *m/z* trendlines for a homologous series, and do not have observable or diagnostic mobility-aligned fragments. Both scenarios would not qualify as a Level 2a under current guidance because a CCS or spectral library match was absent; however, more evidence is available than required for a 2b/c identification. Additionally, IMS libraries are growing, but most spectral libraries do not include CCS values and the vast majority of PFAS in PubChem do not yet have experimental CCS values. Of the 8,099 unique CIDs in PubChem with experimental CCS values, only 208 are PFAS according to the OECD definition implemented in PubChem^5^ (12 Nov. 2024). As spectral and CCS libraries grow and as analytical instrumentation advances, the gray area between a library-based identification and a diagnostic evidence-based identification will likely blur further. Therefore, to align with our goal of reducing ambiguity, we are removing the sublevel distinction in the updated confidence levels. Other key changes from current guidance are described below and summarized in **Table S3.**

**Figure 3.**
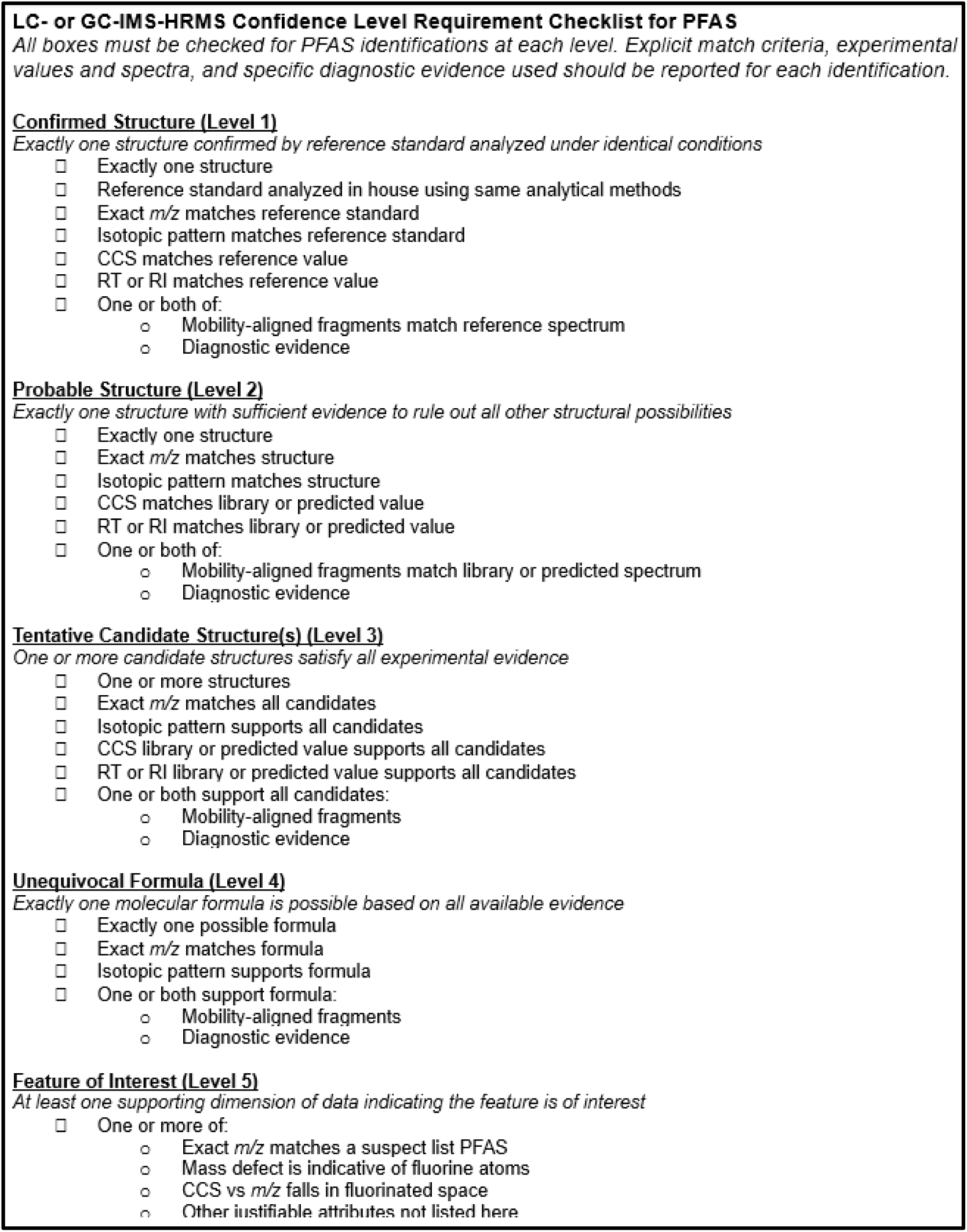
Requirement checklist for PFAS identified using LC- or GC-IMS-HRMS. All listed criteria must be met to assign each confidence level. An extended printable version of the checklist is available in the supplemental information and online.

### Updated PFAS LC- or GC-IMS-HRMS Confidence Levels

The proposed LC- or GC-IMS-HRMS confidence levels for PFAS unify existing guidance from three sources: small molecules confidence levels introduced in 2014,^8^ levels incorporating CCS values from 2020,^12^ and levels specific to PFAS defined in 2022.^3^ The updated levels for PFAS identified using IMS range from 1 to 5 and have no sublevels. These levels also require that acceptable tolerances for all measured dimensions in each experiment must be defined based on the capabilities of their analytical platform. To avoid selecting confidence levels based on desired results, these tolerances should be defined in advance, with rationale clearly described in the experimental methods, and widely accepted as appropriate tolerances for the analytical platform in question. Platform-specific, experiment-appropriate error tolerances will reduce ambiguity and enable uniform confidence level assignment as technology advances. **Table 1** (available as a printable worksheet in **Table S4**) provides suggested tolerances for the Agilent 6560 IMS-QTOF mass spectrometer based on our own experience with PFAS NTA and reference material analysis, widely accepted platform tolerances,^66–69^ and typical workflows used. Future work (beyond the scope of the current manuscript) includes engaging with other IMS PFAS community members to define recommended reporting and tolerances for other instruments and workflows, for broader applicability.

**Table 1.**
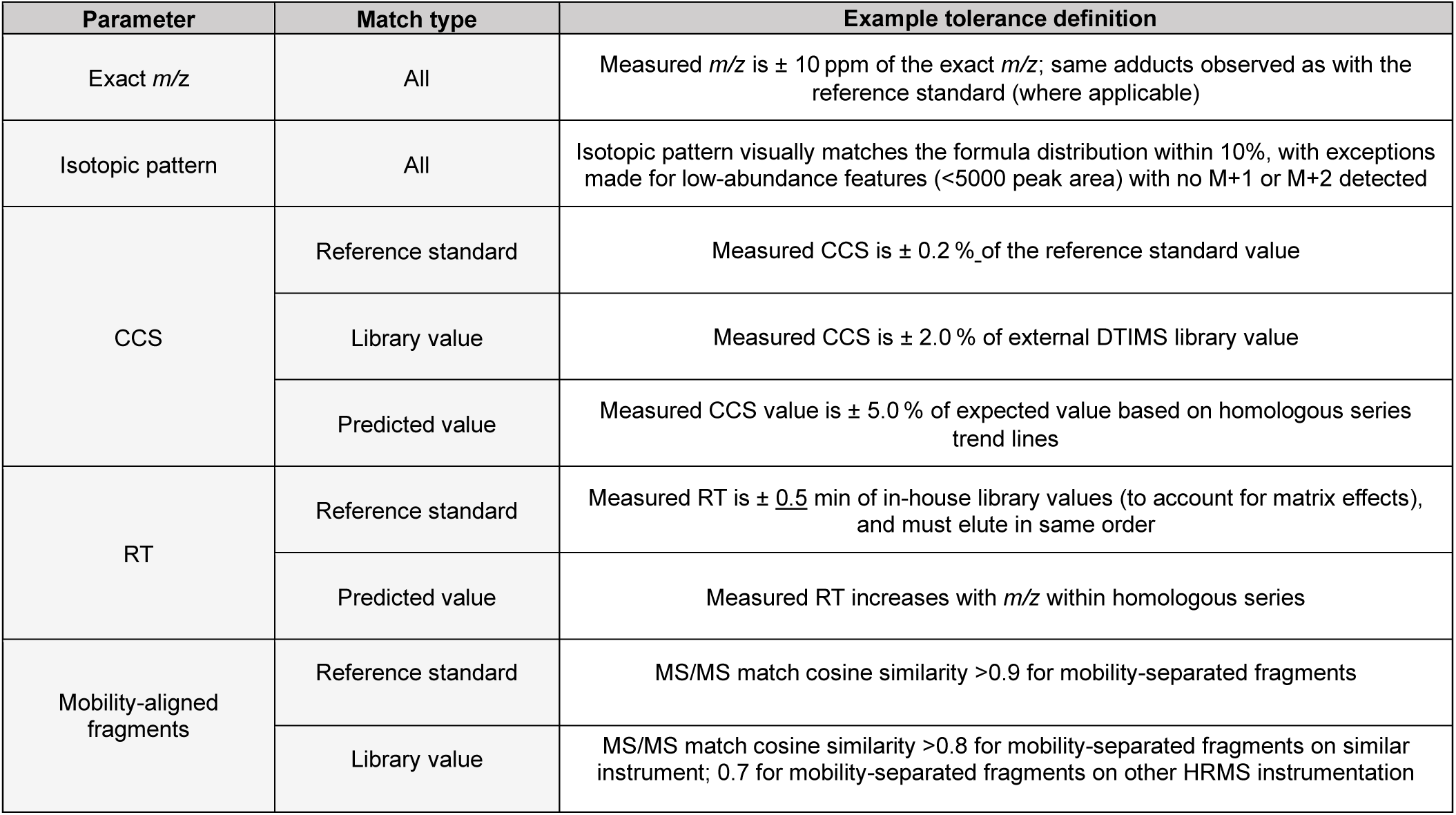
Example tolerance definitions for each measured dimension. These values are recommended default settings for an Agilent 6560 LC-IMS-HRMS platform and have been used for analysis of PFAS in a variety of environmental matrices, in addition to homologous series trendlines for CCS prediction.

| Parameter | Match type | Example tolerance definition |
| --- | --- | --- |
| Exact $m/z$ | All | Measured $m/z$ is $\pm 10$ ppm of the exact $m/z$; same adducts observed as with the reference standard (where applicable) |
| Isotopic pattern | All | Isotopic pattern visually matches the formula distribution within 10%, with exceptions made for low-abundance features (<5000 peak area) with no M+1 or M+2 detected |
| CCS | Reference standard | Measured CCS is $\pm 0.2\%$ of the reference standard value |
| | Library value | Measured CCS is $\pm 2.0\%$ of external DTIMS library value |
| | Predicted value | Measured CCS value is $\pm 5.0\%$ of expected value based on homologous series trend lines |
| RT | Reference standard | Measured RT is $\pm 0.5$ min of in-house library values (to account for matrix effects), and must elute in same order |
| | Predicted value | Measured RT increases with $m/z$ within homologous series |
| Mobility-aligned fragments | Reference standard | MS/MS match cosine similarity >0.9 for mobility-separated fragments |
|  | Library value | MS/MS match cosine similarity >0.8 for mobility-separated fragments on similar instrument; 0.7 for mobility-separated fragments on other HRMS instrumentation |

Broadly, at ***Level 1*,** the identified PFAS is confirmed by matching experimentally validated values from a reference material in all dimensions. At ***Level 2***, sufficient evidence is present that exactly one structure is probable for the identified PFAS, with all other structural possibilities ruled out. At ***Level 3***, at least one candidate structure must fulfill all available evidence, and multiple candidate structures are acceptable. At ***Level 4***, a structure cannot be elucidated but an unequivocal molecular formula is determined, while at ***Level 5*** the feature cannot be identified beyond being a molecule of interest. Definitions and additional requirements are detailed below.

### Definitions

- <u>Match</u>: The measured value for a feature is within the user-defined acceptable error tolerance of the comparison value (*i.e.* the reference standard, library, or predicted value)
- <u>Diagnostic evidence</u>: Experimental observations that corroborate compound identifications. Examples for PFAS could include homologues, compound synthesis information, MS/MS fragments, ionization behavior, parent-transformation product relationships, etc.^8^
- <u>PFAS homologues</u>: Structurally similar compounds differing by one or more repeating structural unit *e.g.* CF_2_, CH_2_CF_2_, or CF_2_O^70^
- <u>Library</u>: A reputable repository containing validated mass spectra and/or CCS values
- <u>Prediction</u>: A calculated spectrum or value obtained from a computational tool or extrapolated from experimental data. Example for PFAS include CCS and RT and values derived from homologous series trendlines, subclass-aligned fragments, and more^3, 23, 70^
- <u>Support:</u> The comparison value aligns with the measured value for a feature and may be outside of the user-defined acceptable error (only applicable at Level 3 and lower).

### Using CCS Values in Confidence Level Assignment

Several different types of IMS separations exist.^17, 71^ Two common methods are drift tube IMS (DTIMS) and traveling wave IMS (TWIMS), both of which have been used to measure CCS values for molecules from a variety of classes, including PFAS.^72^ CCS values are highly reproducible when using the same IMS platform^20, 64, 73^; however, cross-platform comparisons (e.g. comparing CCS values obtained using DTIMS to those obtained from TWIMS) tend to have higher error.^74^ For example, Dodds *et al.* detailed discrepancies between a DTIMS platform and a TWIMS platform, noting CCS measurement differences of around 1% on average between the two platforms, and described a protocol for comparing CCS values across platforms.^72^ Critically, therefore, different tolerances may be required when using library values from a different IMS platform and should be reported transparently. Additionally, some types of IMS separations, such as field asymmetric IMS (FAIMS), do not generate CCS values.^17, 71^ These platforms add an additional separation dimension and may be considered additional diagnostic evidence using our updated guidance; however, this separation is not equivalent to a CCS value.

IMS performance is commonly assessed by evaluating separation efficiency (typically by measuring the resolving power, Rp, calculated using CCS/ΔCCS) and reproducibility (typically with relative standard deviation, or % RSD). Currently, DTIMS platforms can reasonably achieve Rp up to 60 and certain TWIMS platforms can achieve up to 300 without data processing.^72^ Analysts should carefully consider the differences in CCS values for their features of interest and ensure their platform can separate them when defining tolerances. For example, Feature X has a measured CCS of 200.1 Å^2^ using a DTIMS platform with a Rp of 50. Feature X matches two library entries with identical *m/z* and RT values: Isomer A (library CCS value of 199.7 Å^2^) and Isomer B (library CCS value of 200.0 Å^2^). Separating Isomer A and B requires a much higher Rp (> 660) than the DTIMS can reasonably achieve, and it would not be appropriate to annotate Feature X as Isomer B based on having a closer CCS value. Without other evidence, such as diagnostic fragmentation spectra, both Isomer A and Isomer B should both be reported as tentative candidates for Feature X, at a Level 3 confidence as described subsequently, potentially nominating Isomer B as the “most likely” candidate as the CCS value is closer. Certain data processing tools, such as high resolution demultiplexing, may promote higher resolution separations on these platforms and should also be considered when defining tolerances.^75, 76^

### Confirmed Structure (Level 1)

All Level 1 identifications must meet the minimum requirements of exact *m/z*, isotopic envelope, CCS, and RT (in the case of LC) or retention index (RI; in the case of GC) match to a reference standard (**Figure 3**). Reference values from a matching analytical standard must be collected on the same analytical platform as the unknown. These spectra may come from separate injections of the analytical standards or from spiked internal native or mass labeled standards, either analyzed at the same time as the samples of interest, or analyzed previously in-house using identical analytical methods.^8, 12^ The high reproducibility of intra-laboratory CCS values and the ability of IMS to separate PFAS and matrix interferences makes it possible for analysts to use in-house libraries containing RT values, CCS values, and mass spectra without having to run new reference standards alongside experimental samples within the same analytical run. The number of possible Level 1 identifications is thus limited by the number of standards available to the analyst. Existing scales require MS2 spectra for all detected features; however, fragmentation spectra are not always obtainable for low abundance features, which is particularly problematic for environmental analysis of complex biological matrices. Given the high reproducibility of CCS values, the ability of IMS to separate PFAS from interfering biomolecules, and the increased confidence afforded by adding a CCS dimension, mobility-aligned fragments are not required for Level 1 PFAS identifications if diagnostic evidence is available such as the presence of other homologues in the series. Analysts should, however, consider all empirical data available and if anything does not align with the identification, the identification should be downgraded.

### Probable Structure (Level 2)

At Level 2, all evidence in all available dimensions must point to exactly one structure and eliminate all others from contention.^8^ This is the highest confidence level possible when standards cannot be obtained for the molecule in question. All Level 2 identifications must match either a library value or a predicted value (if a library value is not available) for exact *m/z*, isotopic envelope, CCS, and RT (for LC) or RI (for GC). If the measured value does not match the library or predicted value within the predefined, experiment-appropriate tolerance, it is not sufficient to justify a Level 2 identification; however, it may support a Level 3. The library value must come from a reputable spectral or CCS library or literature appropriate to the experiment.^77, 78^ Predicted values may come from a variety of sources, for example CCS vs *m/z* and RT vs *m/z* trendlines for homologous series, and the error of the prediction must be sufficiently narrow to eliminate other candidate structures.^15, 23^ Fragmentation spectra should be used to determine a structure when available; however, it is possible that a combination of MS1 data, CCS values, RT/RI, and diagnostic evidence (such as the presence of homologues at other masses) may be sufficient to justify a Level 2 assignment for some probable structures when no other possibility satisfies all available evidence.

### Tentative Candidates (Level 3)

With Level 3 identifications, at least one candidate structure is supported by all available evidence.^3, 8, 12^ As with Level 2, Level 3 requires either a library or predicted value for exact *m/z*, isotopic envelope, CCS, and RT or RI for all candidate structures. However, at Level 3, multiple tentative candidate structures are permitted, *i.e*., the requirement for exactly one structure only is relaxed. Fragmentation spectra are typically required to determine a structure and should be used when available; however, as with Level 2, diagnostic evidence (see ***Definitions***) may be sufficient to propose a tentative candidate structure in the absence of fragmentation data. When multiple candidates are proposed, it is recommended to report a “most likely” candidate if possible, although we emphasize that the most likely candidate is not equivalent to a Level 2 probable structure. In the earlier example with Feature X (CCS 200.1 Å^2^), Isomer B (CCS 200.0 Å^2^) could be reported as the most likely candidate based on closest library match, with Isomer A (CCS 199.7 Å^2^) also included as a tentative candidate. Various other scoring terms or lines of evidence could be used for selecting a most likely candidate depending on the software approaches used or information available, such as experimental or in silico fragment matching, experimental or predicted CCS values, literature/patent scores, retention information, etc. ^79^Level 3 is also an appropriate assignment when a reference standard or library was used, but the calculated errors are outside of the given tolerance criteria; in this case the measured values do not match the comparison values but may still be used to support a tentative candidate structure. While Metz *et al*.^79^ recently proposed a “probability” approach to rank cases with multiple “n” candidates, this is not yet integrated in this level system while we await community feedback on their proposal.

### Unequivocal Molecular Formula (Level 4)

The definition of a Level 4 identification is an unequivocal formula for the detected feature.^3, 8, 12^ The exact *m/z* and isotopic pattern must match exactly one possible molecular formula. Molecular formula determination typically requires an isotopic envelope and rich fragmentation information, but these are not fixed requirements. The highest scoring formula of a list of possible formulas does not qualify as an unequivocal formula without additional diagnostic evidence, as these are often wrong, especially with the presence of fluorine in the candidate formulas – a fact that is unavoidable in PFAS NTA.

### Feature of Interest (Level 5)

Level 5 is the catchall for any feature in the dataset that cannot be annotated with an unequivocal formula or candidate structure but is still of potential interest to the experimental question.^8, 12^ This does not mean that every remaining feature is annotated with Level 5. Often in environmental NTA there will be thousands of features that do not even qualify for Level 5; for example, matrix lipids when evaluating PFAS, or statistically insignificant features when comparing between two groups of samples (*e.g.,* statistical comparisons of features detected in upstream vs downstream industrial effluent). The requirements for a Level 5 identification include a measured monoisotopic *m/z*, a measured CCS, and at least one piece of evidence that supports the feature being a PFAS of interest. In the move towards automation, “of interest” must be explicitly defined. The additional evidence could be database driven, such as an exact mass match to a suspect list PFAS (equivalent to a Level 5a per the 2022 PFAS levels^3^) with insufficient data to bump up the identification to a Level 4. The evidence could also be experimental (equivalent to a Level 5b per the 2022 PFAS levels^3^), such as a negative mass defect, feature presence in fluorinated space when plotting its CCS vs *m/z*, presence of CF_2_ homologues, presence of CF_2_-containing fragments, or any number of other justifications.

### Discussion and Examples of PFAS Identified with IMS

In typical LC- and GC-HRMS workflows, chromatographic separation and confirmation with diagnostic MS2 fragments is required for compounds with very close *m/z* values. **Figure 4A** shows two PFAS, Hydro-EVE and 6:2 FTS, that have *m/z* values within 10 ppm and require LC separation on most platforms. However, their CCS values are distinct enough that IMS fully resolves these two molecules.^7^ An evaluation of our in-house library^24^ containing experimentally measured RT and CCS values from authenticated standards for 175 PFAS revealed only 3 pairs of PFAS with the potential to overlap with another PFAS in the library when LC, IMS, and HRMS dimensions are all considered with the tolerances shown in **Table 1**, not including branched isomers (*e.g.*, linear PFOA separates from branched PFOA, but individual branched isomers do not always separate from each other).^7^ **Figure 4B** shows one of these pairs, 8:2 FTS and 8:2 FTPA, that may not fully separate using MS1, RT, and CCS depending on the resolving power of the IMS system (remaining pairs shown in **Figure S3**).

**Figure 4.**
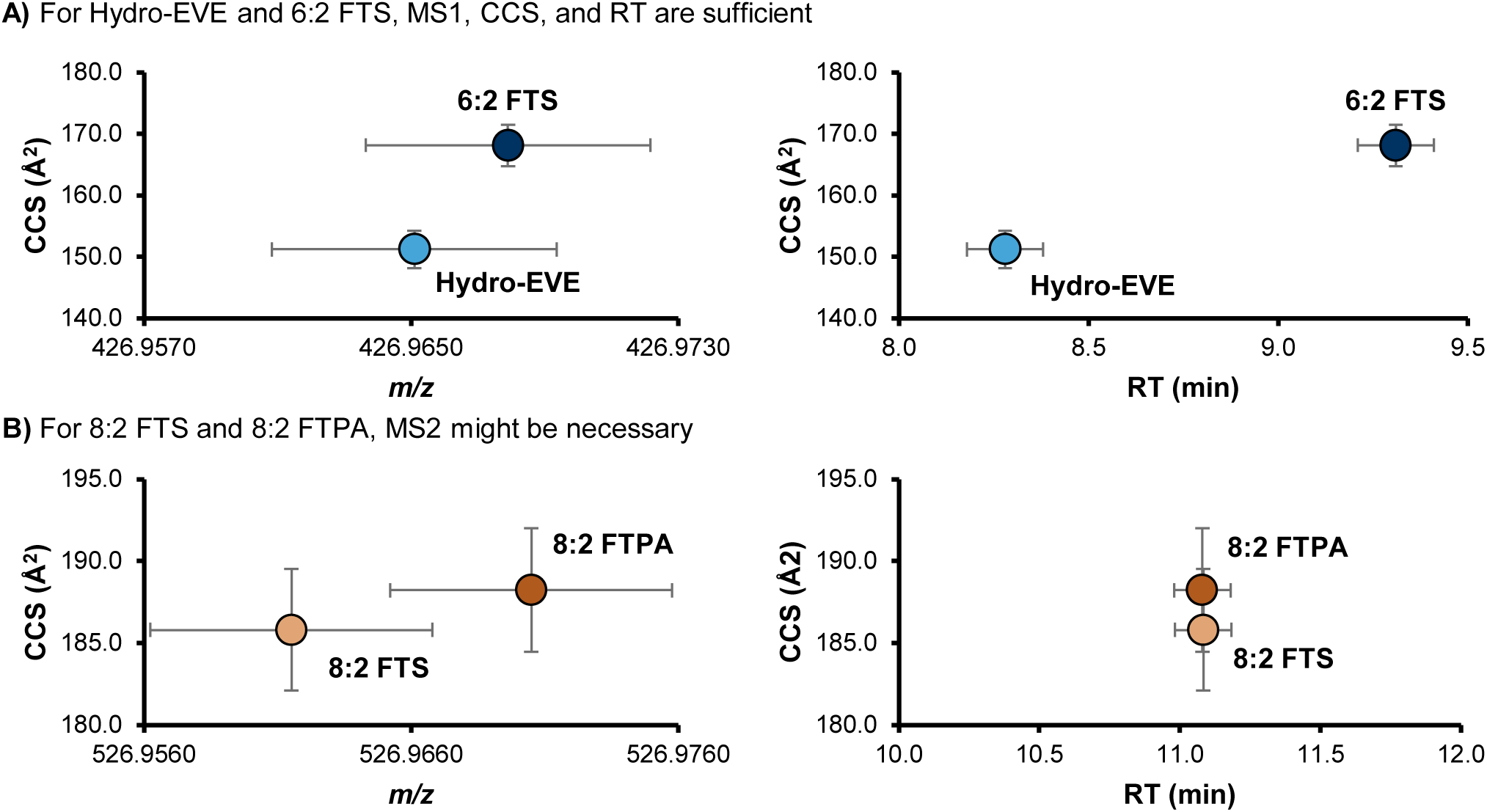
Two examples of PFAS measured using LC-IMS-HRMS. The vertical error bars show ±2.0% CCS, and the horizonal error bars show ± 10 ppm (for *m/z*) and ± 0.1 min (for RT). **A)** The exact *m/z* of deprotonated Hydro-EVE and 6:2 FTS are quite close (426.9651 and 426.9679 respectively, a difference of 6.57 ppm). However, their CCS values (151.2 and 168.1 A^2^) diverge enough (11.1%) to fully resolve these two molecules even if they were to coelute. **B)** On the other hand, 8:2 FTS and 8:2 FTPA have a larger difference in *m/z* (526.9615 and 526.9705, 17 ppm) that are resolved by a tolerance of ± 5 ppm but have a chance of overlapping with a tolerance of ± 10 ppm. Because the CCS values are also close and these molecules coelute in this method, MS2 is recommended to confirm which structures are present in the sample. However, depending on the LC gradient and the resolving power of the IMS platform used, MS2 may not be necessary.

Existing confidence level guidance does not currently consider subjective parameters like geographical knowledge and whether certain PFAS may be coming from manufacturing products in the area. However, such context is often important to the PFAS identification workflow and may sometimes factor into confidence level assignments when assigning tentative Level 3 structures, selecting a most likely candidate at Level 3, bumping a tentative Level 3 candidate to a probable Level 2 structure, or confirming a Level 1 identification in the absence of observable fragments. Our updated guidance permits the analyst to use such knowledge as diagnostic evidence so long as it is reported transparently. For example, if a researcher is studying PFAS in water upstream and downstream of a fluorochemical manufacturer and they have existing knowledge that only one of the candidates is produced by that manufacturer, the user could change the confidence from Level 3 to Level 2. All minimum requirements for mass spectra, CCS, and RT must still be met in this case. Thus, in the example illustrated in **Figure 4**, confidence levels would be assigned as follows, assuming a mass tolerance of ± 10 ppm and a CCS tolerance of ± 0.5%:

- 6:2 FTS and Hydro-EVE: Level 1
- 8:2 FTS and 8:2 FTPA: Level 1 (with diagnostic or fragmentation evidence), Level 3 otherwise (report feature as “8:2 FTS or 8:2 FTPA”)

### Perspective

Effective NTA workflows are essential for monitoring and discovering PFAS and other emerging compounds, and transparent communication of results is critical. Multidimensional techniques like LC-IMS-HRMS are invaluable for analyzing complex environmental and biological samples; however, current methods suffer from inconsistent data quality and confidence reporting. Here, we adapted existing confidence level frameworks to PFAS identified using LC- and GC-IMS-HRMS to improve data quality and reporting of identification confidence in this rapidly growing area of research. Our research experiences amongst study coauthors combined with a dedicated literature search revealed the inconsistent use of existing confidence level guidance for PFAS across a wide range of NTA platforms. To address this gap, we unified and simplified existing confidence level guidance by creating a minimum requirements checklist for PFAS identified at each level. We additionally proposed improvements including user-defined, instrument-specific tolerances for quality control and updated requirements for CCS and MS2 fragmentation across all levels. Recommended tolerances for the platform best known to the authors are given in **Table 1**, and future work will involve consultation with community members to establish other sets of recommended tolerances for other platforms, to ensure broader applicability. These updates provide clear methods for analysts to communicate confidence of PFAS identifications and increase the transparency of NTA results. While this guidance is currently intended for PFAS identified using LC- or GC-IMS-HRMS, researchers using non-IMS platforms focused on molecules other than PFAS may also benefit from these updates. This guidance should be updated as new analytical technologies appear and predictive tools improve, ultimately growing towards automation. To support this, we are developing a web-based application (Small Molecule Identification Scoring Made Easy, or “SMISE”) for automated confidence level assignment as well as additional functionality to aid NTA researchers in proposing and eliminating candidate structures. Release of this tool will enable automated and objective confidence level assignment and unified reporting of evidence used in NTA PFAS identifications.

## Supporting information

Supplemental Checklist

## Declarations

### Data Availability Statement

The Baker lab PFAS LC-IMS-HRMS library is freely available on Zenodo (DOI: https://doi.org/10.5281/zenodo.14341320). A standalone checklist is also available on Zenodo (DOI: https://doi.org/10.5281/zenodo.15441451).

## Acknowledgements

Funding for this work was made possible through grants from the NIH National Institute of Environmental Health Sciences (P42 ES027704, P42ES031007, and T32ES007126), a cooperative agreement with the Environmental Protection Agency (STAR RD 84003201), and support was additionally provided through the Institute for Environmental Health Solutions at the University of North Carolina Gillings School of Global Public Health. ELS acknowledges funding support from the Luxembourg National Research Fund (FNR) for project A18/BM/12341006.

## Supporting Information

The Supplemental Document (PDF) contains supplemental tables (including **S1**: The 38 papers included in the literature analysis. **S2**: Breakdown of experiment types. **S3**: Summary of proposed changes to existing guidance. **S4**: Worksheet for defining tolerances.) and figures (including **S1**: Number of PFAS identified in NTA studies from 2023. **S2**: Examples of inconsistent confidence levels. **S3**: LC-IMS-HRMS library evaluation of overlapping PFAS.).

The Supplemental Checklist (PDF) is a printable version of the checklist shown in **Figure 3**, with definitions added, to serve as a standalone reference

