## Supplemental Checklist for "Updated Guidance for Communicating PFAS Identification Confidence with Ion Mobility Spectrometry"

**LC- or GC-IMS-HRMS Confidence Level Requirement Checklist for PFAS**

*All boxes must be checked for PFAS identifications at each level. Explicit match criteria, tolerances, experimental values and spectra, and specific diagnostic evidence should be reported for each identification.*

**Confirmed Structure (Level 1)**

*Exactly one structure confirmed by reference standard analyzed under identical conditions*

- ☐ Exactly one structure
- ☐ Reference standard analyzed in house using same analytical methods
- ☐ Exact *m/z* matches reference standard
- ☐ Isotopic pattern matches reference standard
- ☐ CCS matches reference value
- ☐ RT or RI matches reference value
- ☐ One or both of:
  - Mobility-aligned fragments match reference spectrum
  - Diagnostic evidence

**Probable Structure (Level 2)**

*Exactly one structure with sufficient evidence to rule out all other structural possibilities*

- ☐ Exactly one structure
- ☐ Exact *m/z* matches structure
- ☐ Isotopic pattern matches structure
- ☐ CCS matches library or predicted value
- ☐ RT or RI matches library or predicted value
- ☐ One or both of:
  - Mobility-aligned fragments match library or predicted spectrum
  - Diagnostic evidence

**Tentative Candidate Structure(s) (Level 3)**

*One or more candidate structures satisfy all experimental evidence*

- ☐ One or more structures
- ☐ Exact *m/z* matches all candidates
- ☐ Isotopic pattern supports all candidates
- ☐ CCS library or predicted value supports all candidates
- ☐ RT or RI library or predicted value supports all candidates
- ☐ One or both support all candidates:
  - Mobility-aligned fragments
  - Diagnostic evidence

**Unequivocal Formula (Level 4)**

*Exactly one molecular formula is possible based on all available evidence*

- ☐ Exactly one possible formula
- ☐ Exact *m/z* matches formula
- ☐ Isotopic pattern supports formula
- ☐ One or both support formula:
  - Mobility-aligned fragments
  - Diagnostic evidence

**Feature of Interest (Level 5)**

*At least one supporting dimension of data indicating the feature is of interest*

- ☐ One or more of:
  - Exact *m/z* matches a suspect list PFAS
  - Mass defect is indicative of fluorine atoms
  - CCS vs *m/z* falls in fluorinated space
  - Other justifiable attributes not listed here

Definitions

- Match: the measured value is within user-defined acceptable error tolerance of a comparison value
- Support: the measured attribute provides evidence that aligns with the proposed structures or formula
- LC: liquid chromatography
- GC: gas chromatography
- IMS: ion mobility spectrometry
- HRMS: high resolution mass spectrometry
- PFAS: per- and polyfluoroalkyl substances
- CCS: collision cross section (obtained from ion mobility separation)
- RT: chromatographic retention time
- RI: chromatographic retention index (typically applicable to GC only)
- Diagnostic evidence: other experimental data supporting the PFAS identification, including but not limited to relevant synthesis information, geographic knowledge, presence of homologues, and parent-transformation product relationships
- Homologue: $CF_2$ or other molecular repeating unit found in known PFAS structures
