## Supplemental Document (PDF) for "Updated Guidance for Communicating PFAS Identification Confidence with Ion Mobility Spectrometry"

Summary: 10 pages, 3 figures, 4 tables

### Contents

Table S1 .......... S3
Table S2 .......... S5
Figure S1 .......... S6
Table S3 .......... S8
Table S4 .......... S9
Figure S3 .......... S10

**Table S1.** The 38 papers on PFAS NTA published in 2023 and included in the in-depth analysis.

| Study ID | PMID | Chromato-graphy used | MS platform used in NTA | IMS used | Matrix category | Experiment category | Confidence levels reported? | Count of PFAS detected with HRMS | 1 | 2 | 2a | 2bc | 3 | 4 | 5 |
|---|---|---|---|---|---|---|---|---|---|---|---|---|---|---|---|
| 3 | 36622151 | LC | Orbitrap | No | Water | PFAS-environmental | Yes-Schymanski | 18 | 0 | 18 | 0 | 18 | 0 | 0 | 0 |
| 28 | 36459821 | GC | TOF | Yes-cIMS | Multiple | PFAS-environmental | No | 25 | na | na | na | na | na | na | na |
| 29 | 36577148 | LC | Orbitrap | No | Standards | PFAS-remediation | No | 52 | na | na | na | na | na | na | na |
| 7 | 36827766 | LC | Orbitrap | No | Water | PFAS-environmental | Yes-Schymanski | 54 | 33 | 16 | na | na | 5 | 0 | 0 |
| 8 | 37058300 | LC | TOF | No | Soil | PFAS-environmental | Yes-Charbonnet | 87 | 0 | 87 | 0 | 87 | 0 | 0 | 0 |
| 9 | 37100129 | LC | TOF | No | Blood | PFAS-environmental | Yes-Schymanski | 93 | 0 | 28 | 0 | 0 | 0 | 0 | 0 |
| 27 | 36787488 | Multiple | Multiple | No | Other industrial | Pollutants-industrial | Yes-Schymanski | 0 | na | na | na | na | na | na | na |
| 6 | 36801580 | LC | Orbitrap | No | Water | PFAS-environmental | Yes-Charbonnet | 30 | 17 | 7 | 1 | 6 | 6 | 0 | 0 |
| 17 | 37648245 | LC | TOF | No | Blood | PFAS-environmental | Yes-Charbonnet | 30 | 13 | 2 | 2 | 0 | 14 | 0 | 1 |
| 30 | 36863195 | GC | Orbitrap | No | Standards | PFAS-other | Yes-other | na | na | na | na | na | na | na | na |
| 31 | 36947591 | GC | Orbitrap | No | Air | PFAS-industrial | No | 5 | na | na | na | na | na | na | na |
| 19 | 37800548 | Multiple | Multiple | No | Multiple | PFAS-environmental | Yes-Schymanski | 82 | 32 | 13 | na | na | 12 | 11 | 0 |
| 20 | 37816443 | LC | Orbitrap | No | Water | PFAS-environmental | Yes-Schymanski | 50 | 0 | 13 | na | na | 27 | 10 | 0 |
| 32 | 37196788 | LC | TOF | No | Water | PFAS-remediation | No | 15 | na | na | na | na | na | na | na |
| 23 | 37923386 | LC | Orbitrap | No | Water | PFAS-environmental | Yes-Schymanski | 44 | 0 | 0 | 0 | 0 | 1 | 0 | 43 |
| 33 | 37207413 | LC | TOF | No | Multiple | PFAS-environmental | No | 2 | na | na | na | na | na | na | na |
| 25 | 37991390 | LC | Orbitrap | No | Soil | PFAS-environmental | Yes-Schymanski | 67 | 47 | 3 | na | na | 15 | 2 | 0 |
| 12 | 37307429 | SFC | TOF | No | Water | Pollutants-environmental | Yes-Schymanski | 3 | 3 | 0 | 0 | 0 | 0 | 0 | 0 |
| 13 | 37394749 | LC | Orbitrap | No | Blood | Pollutants-environmental | Yes-Schymanski | 21 | 4 | 0 | na | na | na | na | na |

| | | | | | | | | | | | | | | | |
|---|---|---|---|---|---|---|---|---|---|---|---|---|---|---|---|
| 14 | 37414869 | LC | Orbitrap | No | Air | Pollutants-environmental | Yes-Schymanski | 2 | 1 | 0 | na | na | 1 | 0 | 0 |
| 1 | 36341487 | LC | TOF | No | Water | PFAS-environmental | Yes-Charbonnet | 47 | 4 | 18 | 12 | 6 | 12 | 4 | 9 |
| 16 | 37590049 | LC | Orbitrap | No | Other industrial | PFAS-industrial | Yes-Charbonnet | 24 | 0 | 0 | 0 | 0 | 6 | 4 | 3 |
| 15 | 37419156 | LC | TOF | No | Other industrial | PFAS-industrial | Yes-Schymanski | 3 | 0 | 3 | 3 | 0 | 0 | 0 | 0 |
| 18 | 37723602 | LC | TOF | No | Blood | PFAS-environmental | Yes-Schymanski | 64 | 16 | 12 | na | na | 3 | 33 | 0 |
| 2 | 36374630 | None | FTICR | No | Water | PFAS-remediation | Yes-Schymanski | 68 | 0 | 5 | na | na | 0 | 63 | 0 |
| 4 | 36722905 | LC | Orbitrap | No | Standards | PFAS-remediation | Yes-Charbonnet | 28 | 0 | 13 | 0 | 13 | 12 | 0 | 1 |
| 34 | 37856848 | LC | Orbitrap | No | Other industrial | PFAS-remediation | No | 37 | na | na | na | na | na | na | na |
| 21 | 37878714 | LC | TOF | Yes-DTIMS | Water | PFAS-environmental | Yes-Charbonnet | 57 | 36 | 8 | 3 | 5 | 3 | 4 | 6 |
| 22 | 37878998 | LC | Orbitrap | No | Water | PFAS-environmental | Yes-Schymanski | 26 | 0 | 26 | 0 | 26 | 0 | 0 | 0 |
| 35 | 37884429 | LC | Not specified | No | Blood | PFAS-environmental | No | 18 | na | na | na | na | na | na | na |
| 10 | 37200532 | LC | TOF | No | Standards | PFAS-remediation | Yes-Charbonnet | 102 | 0 | 33 | 8 | 25 | 11 | 0 | 47 |
| 36 | 37934628 | LC | TOF | No | Other biological | Pollutants-environmental | Yes-other | 3 | na | na | na | na | na | na | na |
| 24 | 37970096 | LC | Orbitrap | No | Multiple | PFAS-industrial | Yes-Schymanski | 22 | 1 | 9 | na | na | 10 | 2 | 0 |
| 37 | 37988936 | GC | TOF | No | Multiple | PFAS-environmental | No | 10 | na | na | na | na | na | na | na |
| 5 | 36736215 | LC | Orbitrap | No | Water | Pollutants-environmental | Yes-Schymanski | 15 | 0 | 0 | 0 | 0 | 1 | 0 | 0 |
| 26 | 38133352 | LC | TOF | No | Multiple | PFAS-environmental | Yes-Schymanski | 3 | 1 | 0 | 0 | 0 | 0 | 0 | 1 |
| 11 | 37222023 | LC | Orbitrap | No | Blood | Pollutants-environmental | Yes-Schymanski | 11 | 0 | 3 | 0 | 0 | 0 | 0 | 0 |
| 38 | 37503704 | Multiple | TOF | No | Other industrial | PFAS-industrial | No | 0 | na | na | na | na | na | na | na |

**Table S2.** Breakdown of types of experiments from 38 papers on PFAS NTA published in 2023.

<table>
<tr><th colspan="6">Analytical platform</th></tr>
<tr><th>HRMS instrument</th><th>Count</th><th>Chromatography</th><th>Count</th><th>IMS</th><th>Count</th></tr>
<tr><td>Orbitrap</td><td>18</td><td>LC</td><td>29</td><td>None</td><td>36</td></tr>
<tr><td>TOF</td><td>16</td><td>GC</td><td>4</td><td>cIMS</td><td>1</td></tr>
<tr><td>Multiple</td><td>2</td><td>Multiple</td><td>3</td><td>DTIMS</td><td>1</td></tr>
<tr><td>FTICR</td><td>1</td><td>SFC</td><td>1</td><td></td><td></td></tr>
<tr><td>Not specified</td><td>1</td><td>None</td><td>1</td><td></td><td></td></tr>
<tr><th colspan="6">Experiment type</th></tr>
<tr><th colspan="2">Molecules analyzed</th><th colspan="4">Matrix analyzed</th></tr>
<tr><th>PFAS specific</th><th>Count</th><th>Matrix category</th><th>Count</th><th>Water type</th><th>Count</th></tr>
<tr><td>Environmental</td><td>19</td><td>Water</td><td>12</td><td>Wastewater</td><td>5</td></tr>
<tr><td>Remediation</td><td>6</td><td>Blood</td><td>6</td><td>Ground water</td><td>4</td></tr>
<tr><td>Industrial</td><td>5</td><td>Multiple</td><td>6</td><td>Surface water</td><td>3</td></tr>
<tr><td>Other</td><td>1</td><td>Other</td><td>6</td><td>Drinking water</td><td>1</td></tr>
<tr><td>Pollutants (including PFAS)</td><td>Count</td><td>Standards</td><td>4</td><td>Sea water</td><td>1</td></tr>
<tr><td>Environmental</td><td>6</td><td>Soil</td><td>2</td><td>Passive samplers</td><td>1</td></tr>
<tr><td>Industrial</td><td>1</td><td>Air</td><td>2</td><td></td><td></td></tr>
</table>

**Definitions**
HRMS: high resolution mass spectrometry
TOF: time of flight
FTICR: Fourier transform ion cyclotron resonance
LC: liquid chromatography
GC: gas chromatography
SFC: supercritical fluid chromatography
IMS: ion mobility spectrometry
cIMS: cyclic IMS
DTIMS: drift tube IMS

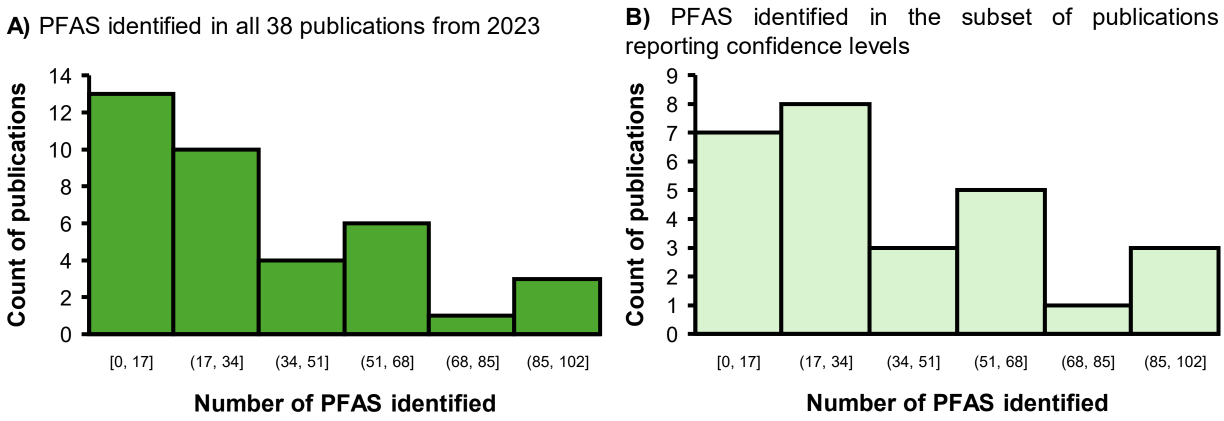


**Figure S1**. **A)** Number of PFAS identified in papers on PFAS NTA published in 2023 in 37 of the 38 papers evaluated (one of the 38 papers evaluated did not report a number of PFAS identified and is not included here). Min = 0, max = 102, median = 25, average = 32.3 PFAS identified per study. **B)** The subset of 27 papers that also reported a confidence level on a 1-5 scale. Min = 0, max = 102, median = 30, average = 38.1 PFAS identified per study.

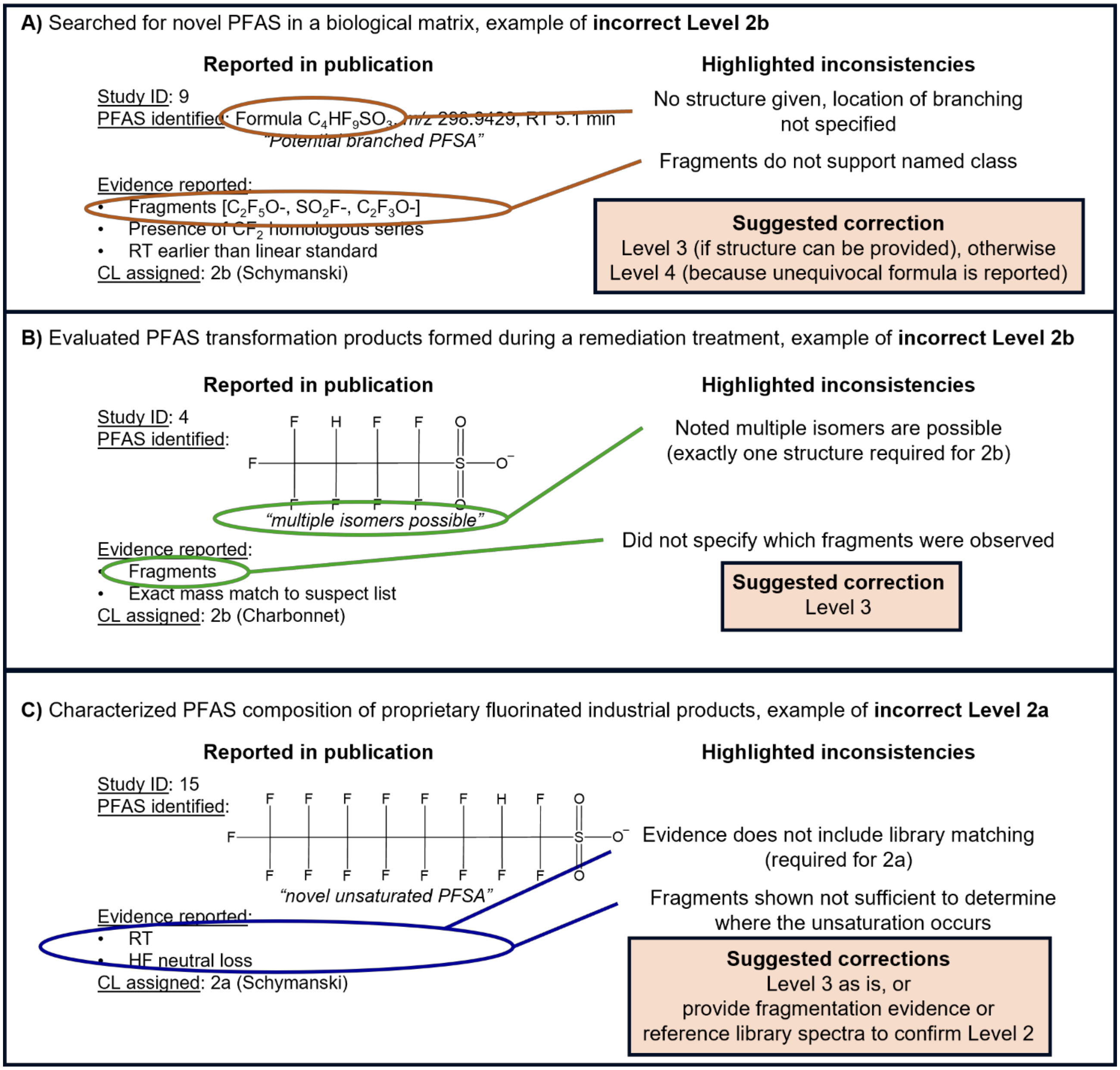


**Figure S2.** Examples of confidence levels assigned in a manner inconsistent with the scale cited by each study. **A)** This study cited the Schymanski scale and assigned a Level 2b to a molecule reported as a "potential branched PFSA"; however, no structure was provided and the reported fragments do not match with the implied structure for a branched PFSA (a fully fluorinated carbon chain with a sulfonic acid head group). **B)** This study cited the Charbonnet confidence scale, assigned a Level 2b to a structure for an unsaturated PFSA, and noted "multiple isomers possible". Fragmentation evidence could potentially distinguish between different isomers; however, if multiple isomers were observed it would be better to report all of them individually. **C)** This study cited the Schymanski scale and assigned a Level 2a to a structure referred to in the text as a "novel unsaturated PFSA" (author note: this could also be termed hydrogen substituted PFSA, but the name used in the original study has been retained for consistency). Level 2a requires spectral matching to a library and therefore it is highly unlikely that truly novel (i.e. previously undiscovered) structures could be elucidated in this manner. Insufficient evidence is provided to support a Level 2 assignment.

**Table S3**. Summary of proposed changes to confidence level guidance for PFAS identifications made using LC-IMS-HRMS

| Attribute | Current guidance | Proposed change |
|---|---|---|
| Mobility-aligned fragments | $MS^n$ "matching" library or reference spectra,[1, 2] minimum 1, 2, or 3 diagnostic or subclass-aligned fragments[3] | Not required when diagnostic evidence is available |
| CCS value | Must match library CCS for Level 2a or predicted CCS for Level 2b or 3[2] | Library or predicted value is acceptable if within user-defined tolerance |
| RT value | Must match reference standard ≤ 0.1 min for Level 1, must match library for Level 2a[2]<br>Not required for homologous series at Level 2b or 3[3] | Library or predicted RT or RI value is acceptable if within user-defined tolerance |
| PFAS-specific sublevels | Confidence decreases in order: 1a > 1b > 2a > 2b and 2c > 3a > 3b > 3c and 3d > 4 > 5a and 5b[3] | Levels 1-5 only, no sublevels |

Table References

(1) Schymanski, E. L.; Jeon, J.; Gulde, R.; Fenner, K.; Ruff, M.; Singer, H. P.; Hollender, J. Identifying Small Molecules via High Resolution Mass Spectrometry: Communicating Confidence. Environmental Science & Technology 2014, 48 (4), 2097-2098. DOI: 10.1021/es5002105.

(2) Celma, A.; Sancho, J. V.; Schymanski, E. L.; Fabregat-Safont, D.; Ibáñez, M.; Goshawk, J.; Barknowitz, G.; Hernández, F.; Bijlsma, L. Improving Target and Suspect Screening High-Resolution Mass Spectrometry Workflows in Environmental Analysis by Ion Mobility Separation. Environmental Science & Technology 2020, 54 (23), 15120-15131. DOI: 10.1021/acs.est.0c05713.

(3) Charbonnet, J. A.; McDonough, C. A.; Xiao, F.; Schwichtenberg, T.; Cao, D.; Kaserzon, S.; Thomas, K. V.; Dewapriya, P.; Place, B. J.; Schymanski, E. L.; et al. Communicating Confidence of Per- and Polyfluoroalkyl Substance Identification via High-Resolution Mass Spectrometry. Environmental Science & Technology Letters 2022, 9 (6), 473-481. DOI: 10.1021/acs.estlett.2c00206.

**Table S4**. Worksheet for defining platform-specific tolerances of each measured dimension. Suggested definitions are provided and users should modify the suggestions as appropriate.

| Parameter | Match type | Tolerance definition |
|---|---|---|
| Exact *m/z* | All | Measured *m/z* is ± _________ of the exact *m/z*; same adducts observed as with the reference standard |
| Isotopic pattern | All | |
| CCS | Reference standard | Measured CCS is ± _____ of the reference standard value / in-house library value |
| | Library value | Measured CCS is ± _____ of external library value |
| | Predicted value | |
| RT or RI | Reference standard | Measured RT / RI is ± ______ of internal / external standards<br>Measured RT / RI is ± ______ of in-house library value from identical chromatographic method |
| | Library value | Measured RT is ± ______ of library value and must elute in the same order as other detected library matches |
| | Predicted value | |
| Mobility-aligned fragments | Reference standard | MS/MS match cosine similarity is > ________ |
| | Library value | Same instrument: MS/MS match cosine similarity is > ________<br>Different instrument: MS/MS match cosine similarity is > ________ |

**A)** The 6 overlapping Baker Group library entries using ± 20 ppm *m/z* difference, ± 5 min RT difference, and ± 5.0% CCS.

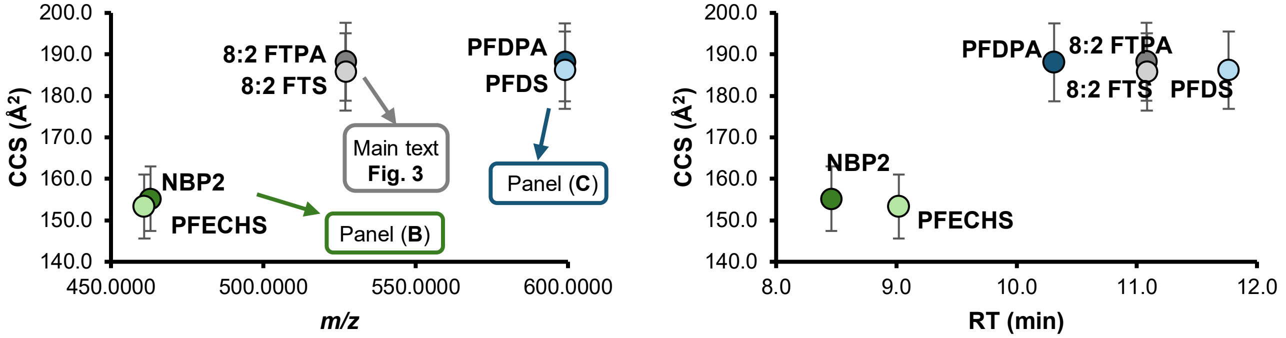


**B)** NBP2 and PFECHS using tolerances of ± 10 ppm, ± 0.1 min, and ± 0.2% CCS.

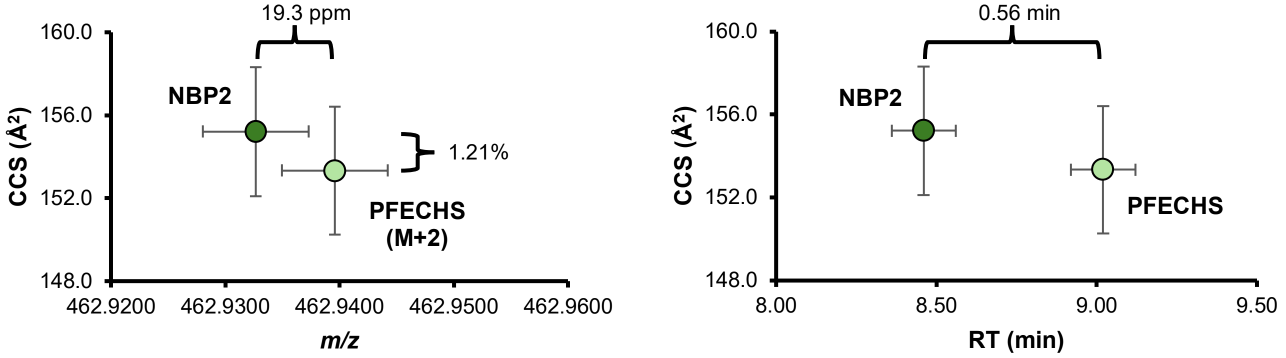


**C)** PFDPA and PFDS using tolerances of ± 10 ppm, ± 0.1 min, and ± 0.2% CCS.

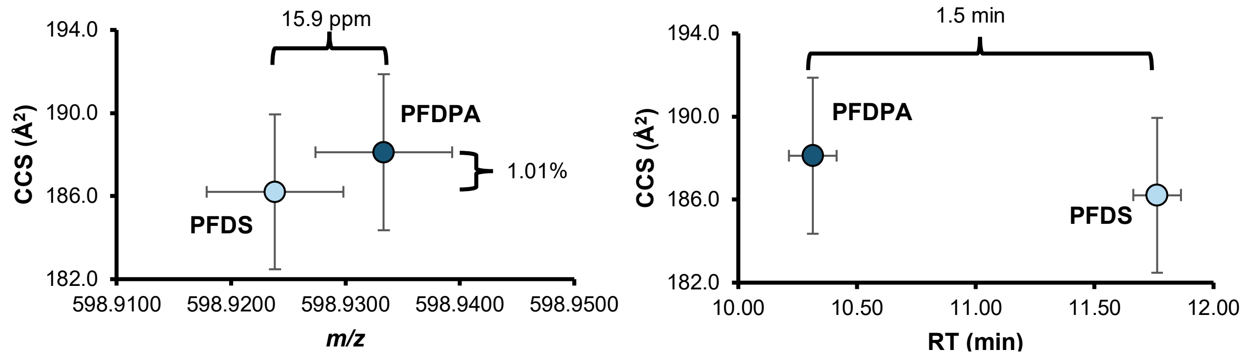


**Figure S3.** Evaluation of our LC-IMS-HRMS library revealed only 6 PFAS with the potential to overlap with another PFAS when considering *m/z* and CCS. CCS and RT values are from the Baker Group PFAS LC-IMS-MS library, v3, available on Zenodo at DOI: https://doi.org/10.5281/zenodo.13387492. **A)** All 6 PFAS in the library found to overlap (excluding branched isomers of PFOA and PFOS, which have been described elsewhere) using wide tolerances of ± 20 ppm *m/z* difference and +1 or +2 Da to account for isotopic envelopes, ± 5 min RT difference, and ± 5.0% CCS. **B)** Nafion Byproduct 2 (NBP2) and PFECHS are 19.3 ppm different and should typically resolve with HRMS alone. However, care should be taken especially if either peak is particularly abundant, because these molecules elute closely and their CCS values are also quite close (1.21%). MS2 may be advised for confirming these structures in some cases, depending on the tolerance of the instrument. **C)** PFDS and PFDPA are 15.9 ppm different in *m/z*, 1.01% different in CCS, and elute 1.5 min apart on our typical LC gradient. If using a different gradient or if matrix effects are observed, MS2 is recommended to confirm which structures are present in the sample.
